# RNABSdb and 3plex enable deep computational investigation of triplex forming lncRNAs

**DOI:** 10.1101/2022.07.06.496678

**Authors:** Chiara Cicconetti, Andrea Lauria, Valentina Proserpio, Annalaura Tamburrini, Mara Maldotti, Salvatore Oliviero, Ivan Molineris

## Abstract

Long non-coding RNAs (lncRNAs) regulate gene expression through different molecular mechanisms, including DNA binding. We curated the first database of RNA Binding Sites (RNABSdb) by harmonising publicly available raw-data of RNA-DNA binding experiments. This resource is crucial to enable systematic studies on transcriptional regulation driven by lncRNAs. Focusing on high quality experiments, we find that the number of binding sites for each lncRNAs varies from hundreds to tens of thousands. Despite being poorly characterised, the formation of RNA:DNA:DNA triple helices (TPXs) is one of the molecular mechanisms that allows lncRNAs to bind the genome and regulate gene expression. We developed 3plex, a software able to predict TPXs *in silico*. We show that 3plex outperforms previous existing approaches leveraging the data collected in RNABSdb for lncRNAs known to form functional TPXs. Moreover this analysis shows that TPXs tend to be shorter and more degenerated than previously expected. Finally, we applied 3plex to all the lncRNAs collected in RNABSdb and we show that the majority of them could directly bind the genome by TPXs formation.

Data and software are available at https://molinerislab.github.io/RNABSdb/ and https://github.com/molinerisLab/3plex.

## 2. Introduction

When pervasive transcription of mammalian genomes was first discovered (1), the prevailing hypothesis was that long non protein-coding RNAs were just “junk RNAs” resulting from random and unregulated RNA polymerase II activity. This theory was supported by RNA expression studies on bulk tissues, which showed that lncRNAs are expressed at lower levels compared to mRNAs. It was unclear whether this was caused by a uniform low expression level of lncRNAs across the cells of a tissue, or because lncRNAs are expressed at high level only in specific cellular subpopulations. This latter hypothesis also suggested that lncRNAs, previously thought to be transcriptional noise, might have highly specialised roles in specific cell types. Thanks to single-cell transcriptomics, an increasing amount of evidence in different biological contexts is supporting this second hypothesis (2–5) and several lncRNAs are reported to be tissue or cell-line specific (6).

Nowadays, the “junk RNA” hypothesis is almost completely discarded, and many lncRNAs have been shown to regulate gene expression both transcriptionally and post-transcriptionally (7). Focusing on transcriptional regulation, key tools to investigate the functional role of lncRNAs are represented by genome wide RNA-DNA binding assays. Several different techniques exist that enable the identification of the genomic RNA binding regions (listed in Table 1). Amongst those, the most used is Chromatin isolation by RNA purification (ChIRP-seq). In this paper we will generically refer to any of these assays as RNABS-seq (RNA Binding Sites Sequencing).

**Table 1.** List of experimental methods used to isolate RNA genomic interaction sites.

| Acronym | Name | Method Description | RNA specificity | DOI (reference) |
| --- | --- | --- | --- | --- |
| ChIRP-seq | Chromatin isolation by RNA purification | Hybridization of biotin-labelled antisense oligos to target RNA | specific for a single RNA | 10.3791/3912 (38) |
| CHART-seq | Capture Hybridization Analysis of RNA Targets | Hybridization of biotin-labelled antisense oligos to target RNA | specific for a single RNA | 10.1002/0471142727.mb2125s101 (39) |
| RAP-seq | RNA affinity purification | Hybridization of biotin-labelled antisense oligos to target RNA | specific for a single RNA | 10.1126/science.1237973 (40) |
| ChOP-seq | Chromatin Oligo affinity Precipitation | Hybridization of biotin-labelled antisense oligos to target RNA | specific for a single RNA | 10.1038/ncomms8743 (35) |
| ASO-Capture-Seq | Anti-Sense Oligonucleotide mediated capture | Hybridization of biotin-labelled antisense oligos to target RNA (deproteinized chromatin) | specific for a single RNA | 10.1093/nar/gky1305 (41) |
| TRIP-seq | Triple helix-sequencing assay | Enrich triplex-forming sequences relative to nucleosome positions. | all genomic regions involved in triplexes (no RNA specificity) | 10.1016/j.molcel.2019.01.007 (42) |
| MARGI-seq | Mapping RNA-genome interactions | Isolation of genome-wide RNA-chromatin interactions through proximity ligation | all RNAs | 10.1016/j.cub.2017.01.011 (43) |
| GRID-seq | Global RNA Interactions with DNA | Isolation of genome-wide RNA–chromatin interactions through proximity ligation | all RNAs | 10.1038/nbt.3968 (44) |
| RADICL-seq | RNA And DNA Interacting Complexes Ligated | Isolation of genome-wide RNA–chromatin interactions through proximity ligation | all RNAs | 10.1038/s41467-020-14337-6 (45) |
| ChAR-seq | Chromatin-Associated RNA sequencing | Isolation of genome-wide RNA–chromatin interactions through proximity ligation | all RNAs | 10.7554/eLife.27024 (46) |

Despite the amount of data produced, we are still missing a curated collection of publicly available RNABS-seq data that would be instrumental to enable systematic studies on transcriptional regulation driven by lncRNAs. Thus we created the RNA Binding Sites database (RNABSdb), a collection of all the publicly available genome wide RNA binding experiments, uniformly re-analysed with the ChIP-seq ENCODE pipeline.

One of the molecular mechanisms that enables lncRNAs to bind specific DNA sequences *in cis* or *in trans* is the formation of RNA:DNA:DNA triple helices (See Fig 1B). These structures are formed by an RNA molecule that, laying on the DNA major groove, establishes hydrogen bonds with DNA purine nucleotides already paired in the double helix. Those RNA-DNA hydrogen bonds do not follow the usual Watson-Crick laws, but the Hoogsteen rules (8,9). The bound portion of the lncRNA is named triplex forming oligo (TFO) and the cognate portion in the DNA molecule is named triplex target site (TTS).

**Figure 1.**
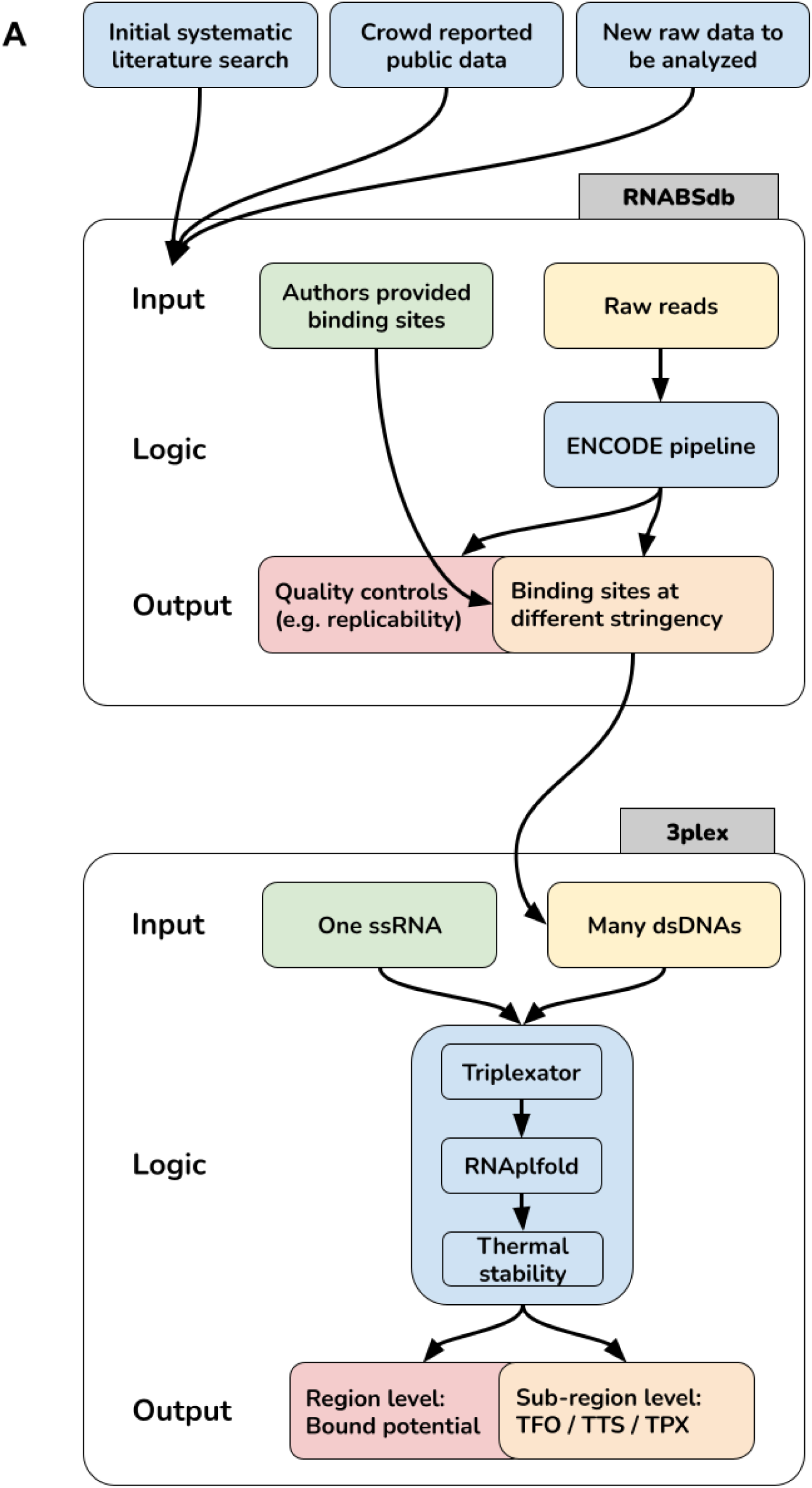
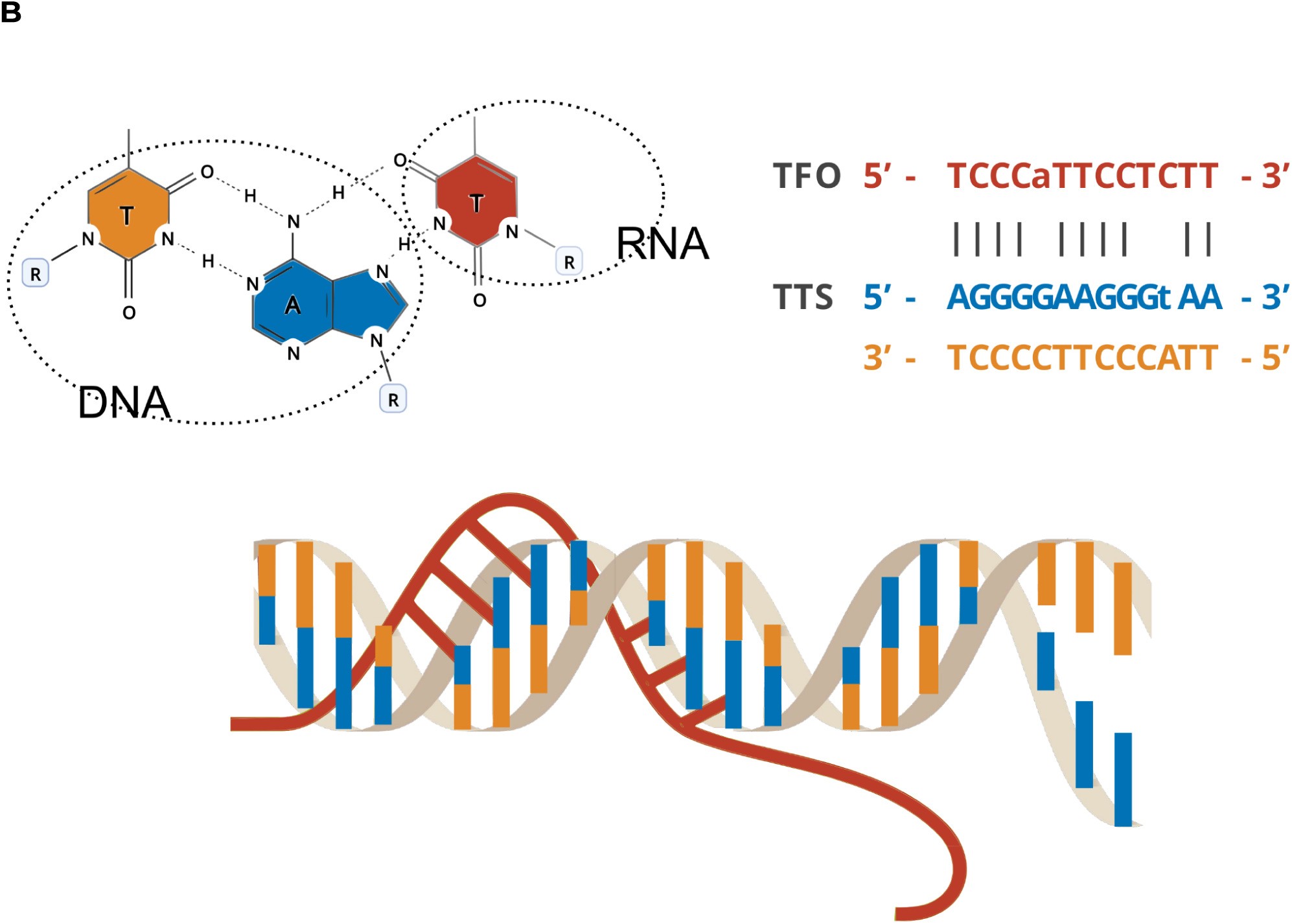
(**A**) Schematic representation of our workflow: we collected publicly available RNA-DNA interaction data in a new database, RNABSdb. We uniformly annotated and processed these data using the ENCODE ChIP-seq pipeline, obtaining quality controls and binding sites at different stringency. This consistent collection was used to test our TPX prediction software (3plex) that integrates methods from Triplexator, RNAplfold and LongTarget. (**B**) Graphic illustration showing an example of Hoogsteen hydrogen bond and a RNA:DNA:DNA triple helix. The sequence alignment shows an example of a TPX involving a parallel oriented pyrimidine TFO (red). The blue and orange strings represent the interacting TTS located on the DNA. Each grey line represents a Hoogsteen pairing (two hydrogen bonds). Lowercase letters represent nucleotides in disagreement with Hoogsteen rules.

Some lncRNAs are known to regulate gene expression of specific target genes through binding to specific TTS in their DNA regulatory regions, acting analogously to transcription factors. For example, the murine lncRNA EPR binds on Arrdc3 promoter, thus activating its expression and modulating the epithelial to mesenchymal transition (10). Similarly, LncSmad7 binds and recruits p300 to enhancer regions *in trans*, triggering their acetylation and transcriptional activation of their target genes, controlling the expression of key stemness regulators (11). Other examples can be found in recent reviews (12,13).

Nowadays, there are few bioinformatic software available to predict the ability of a lncRNA to form triplexes that take in account the ability to form TFO and TTS and the pairing with Hogsteen rules.

Triplex Domain Finder is one of the most used (14), and it is based on the logic of the Triplexator algorithm (15). Triplexator has many parameters, one of the most relevant is the minimum length allowed for TPX, that is set by default at 16nt and cannot be lowered below 10 without modifying the source code. Moreover this parameter is often increased in most of the applications. As an example, Jalaly et al. (16) performed a genomic survey of potential triplexes in the human genome by using a cutoff of 35 for the TPX length. LongTarget is an alternative to Triplexator that focuses on even longer TPX (17) (see Table 2).

**Table 2.** comparison of TPX prediction software.

| Name | Easy installation using containers | Computation time | Minimal - default TPX length | Thermal stability computation | Secondary structure integration | Last update on github |
| --- | --- | --- | --- | --- | --- | --- |
| **3plex** | yes | low | 6-10 | yes | yes | 2022 |
| **Triplexator** | no | low | 10-16 | no | no | 2013 |
| **LongTarget** | no | high | NA-20 | yes | no | 2018 |
| **TDF** | no | low | 10-16 | no | no | 2021 |

There are two principal reasons to focus on long TPX: 1) the identification of small TPX is computationally demanding, 2) longer sequences are required for a stronger and more specific match. For these reasons, shorter TPX have been so far mainly neglected in this type of analysis.

To better characterise the TPX mechanism, we developed 3plex, a TPX prediction software, by integrating Triplexator rules, computational evaluation of thermal stability and RNA secondary structure prediction.

Using data included in RNABSdb as positive control, we evaluated the performance of 3plex and showed that it outperforms the state of the art in this field. We found that lncRNAs tend to bind DNA with short and degenerated TPXs and we revealed that many lncRNAs not previously reported to form triplexes could directly bind the DNA with this mechanism.

## 3. Methods

### 3.1. Collection of high throughput RNA:DNA interaction data

We collected high-throughput RNA:DNA interaction data primarily through manual mining of literature published between November 2011 (when the first ChIRP-seq was published) and February 2022. Gene Expression Omnibus, PubMed and European Nucleotide Archive databases were queried using as keywords the experimental techniques that map RNA-binding sites at the whole genome level including:ChIRP-seq, CHART-seq, ChOP-seq, RAP-seq, MARGI-seq, GRID-seq and RADICL-seq. RNA molecules were uniformly annotated using GENCODE gene symbols. We excluded from the analysis experiments without available raw data or with unclear annotation of the replicates. Xist has been excluded because the experimental data were limited to X chromosome (18).

The data related to some recent papers have not been analysed because published after the data collection was completed (19,20), they will be included in upcoming RNABSdb releases.

### 3.2. ENCODE pipeline on RNABS-seq data

We downloaded raw fastq from GEO or ENA and analysed them using the ENCODE transcription factor pipeline (https://www.encodeproject.org/chip-seq/transcription_factor/ version v1.6.1) (21).

In the ENCODE pipeline, after SPP peak calling (22) with low stringency, replicate concordance is measured by calculating peaks Overlap or IDR (23). Some of the experiments didn’t pass the IDR filter on both the Rescue Ratio and the Self Consistency Ratio, and in these cases no IDR consistent peaks were found. The pipeline also returns a less stringent peak selection without applying IDR procedure, this set is named “Overlap”. We further filtered the “Overlap” peaks sets considering only those having a SPP q-value below 0.05.

Apart from IDR (conservative and optimal) and Overlap (conservative and optimal) peaks, we created a selection of top 1000 peaks obtained as follows: 1) preliminary selection of IDR conservative peaks, 2) if the number of selected peaks is below 1000, inclusion of the best IDR optimal peaks, not overlapping with IDR conservative ones, 3) if the number of selected peaks is still below 1000, addition of the best Overlap conservative peaks that do not overlap with any of the IDR peaks.

#### 3.2.1. Replicate handling

The experimental designs were different across data collected. The majority of the experiments used a couple of even/odd sets of probes that we defined as replicate 1 and 2 in the ENCODE pipeline.

ENCODE guidelines suggest to evaluate the reproducibility of the data by computing:

- N1: Replicate 1 self-consistent peaks (comparing two pseudoreplicates generated by subsampling Rep1 reads)
- N2: Replicate 2 self-consistent peaks (comparing two pseudoreplicates generated by subsampling Rep2 reads)
- Nt: True Replicate consistent peaks (comparing true replicates Rep1 vs Rep2)
- Np: Pooled-pseudoreplicate consistent peaks (comparing two pseudoreplicates generated by subsampling pooled reads from Rep1 and Rep2)
- Self-consistency Ratio: max(N1,N2) / min(N1,N2)
- Rescue Ratio: max(Np,Nt) / min(Np,Nt)

Then the reproducibility test returns “ideal” if Self-consistency Ratio >2 and Rescue Ratio > 2, “acceptable” if one of the two ratios is <2, “concerning” if both the ratios are < 2.

#### 3.2.2. Correlation between number of peaks and QC parameters

We used the Spearman coefficient (cor.test in R) to evaluate the correlations between the number of called peaks and QC parameters reported by the ENCODE pipeline over each experiment analysed. For QC parameters having more than one value per experiment (e.g. one value for each replicate), we aggregated the values considering the average, minimum and maximum values independently before the correlation.

For the IDR conservative number of peaks we found a significant correlation with the Rescue Ratio: this is by construction, because the number of peaks is directly indicated in the Rescue Ratio formula. None of the other QC parameters correlates significantly with IDR conservative number of peaks.

### 3.3. 3plex scoring method based on thermal stability

Given a TPX found according to constraint parameters of Triplexator, a triplet is defined as a group of two Watson-Crick interacting nucleotides in the dsDNA and one Hoogsteen interacting nucleotide in the ssRNA. 3plex implements a new scoring method based on experimentally determined triplets stability data derived from He et al. (17).

To compute the normalised stability, that is the length normalised by the average thermal stability of TPXs on the entire dsDNA considered, we used bedtools merge to avoid overestimation of stability. Since different TPX can overlap, for each set of overlapping TPX we considered only the one with maximal stability. Hence, we selected the maximal stability value among all merged TTSs for each dsDNA and we divided it by the dsDNA length.

This scoring method undercounts the stability for each dsDNA, indeed the contribution to the stability of some dsRNA portion is discarded because of partial overlap. Using RNABSdb binding sites of 7 lncRNAs with experimentally validated functional TPX, we compared the performance of this scoring in terms of AUC with an alternative that overcounts the stability by ignoring overlaps. We found that the undercounting performs similarly or slightly better than the overcounting.

### 3.4. RNA secondary structure prediction

RNAplfold from the ViennaRNA package (24) used with *-u* option measures the probability that stretches of *n* sequential nucleotides on the RNA sequence are unpaired. As suggested by the authors, this option allows to compute the probability that a stretch of n consecutive nucleotides is unpaired, which is useful for predicting possible binding sites. Given an RNA sequence, we associated to each nucleotide the probability that a window of length 8 centred on that position is single stranded. Subsequently, we masked nucleotides from the RNA sequence which report a probability value lower than a certain cutoff. The selected cutoffs enable masking the 10%, the 20% or the 50% of the sequence (ss10, ss20, ss50). The unmasked sequence is defined as ss0.

### 3.5. Positive and negative regions for parameter evaluation and AUC computation

Positive regions are peaks reported in RNABSdb, considering the 5 different filtering methods available as described in section 3.1. If more than one even/odd set were provided, the intersection of the called peaks is considered as a positive set for 3plex evaluation.

Negative regions are randomly selected from the genome maintaining the same length distribution of the positive ones using bedtools shuffle. Positive regions, genome gaps and ENCODE blacklist regions are excluded from the selection.

ROC curves and AUC are obtained for different scoring methods using pROC package version 1.17.0.1 (25). We compared different AUC using roc.test of the same package (two sided Delong’s test).

One sided Mann-Whitney test was performed using the R function wilcox.test in order to test the significance of the difference between the scores of the positive and negative regions.

Benjamini-Hochberg correction on p-values was computed using the R function p.adjust setting method to “BH”.

### 3.6. Linear models for parameters relevance evaluation

To evaluate the relevance of parameters in the TPX prediction, we explored the parameters space of RNABSdb and 3plex. The complete list of parameters evaluated is

- Peak filtering method, described in section 3.1 and 3.2.1, with possible values:
  - IDR conservative
  - IDR optimal
  - Overlap conservative, filtered at q<0.05
  - Top1000 as described in section 3.2
  - The set of peak provided by the authors of the experiment
- ssRNA, a factorial variable indicating the specific lncRNA,
- the number of peaks in RNABSdb for the given lncRNA and peak filtering method,
- single strandedness, as described in section 3.4, with possible values:
  - ss0,
  - ss10,
  - ss20,
  - ss50,
- the minimal length of TPX to be considered, with possible values:
  - 16
  - 12
  - 10
  - 8
- the minimal error rate allowed in a TPX, with possible values:
  - 10%
  - 20%
- the maximum number of consecutive error allowed in TPX, with possible values:
  - 1
  - 3
- the minimal guanine rate allowed in TTS, with possible values:
  - 10%
  - 40%
  - 70%
- repeat filter, indicating if low complexity regions in ssRNA should be masked or not
- the triplex scoring method as described in section 3.3.2 and 4.3.2, with possible values:
  - Triplexator potential
  - Normalised stability
  - Triplexator best score
  - Best stability

We considered 2 different models.

The first model aims to investigate the effect of minimal length, keeping the Triplexator parameters fixed at the value suggested by Matveishina et al. (26) (namely: minimal length=10, error rate=20%, guanine rate=40%) or the Triplexator default, we also excluded 3plex specific parameters (thus excluding thermal stability triplex related scoring methods and secondary single strandedness). We computed AUC for 7 lncRNAs having experimental evidence of functional TPX in each configuration of parameters (namely peak filtering method and minimal length), then we fitted a linear model (lm function on R) to study the dependency of AUC on different parameters controlling for the ssRNA as covariate (formula: AUC ∼ peak filtering method + ssRNA + n peaks + minimal length) model. We also fitted an analogous nested model with the same formula, but excluding the minimal length parameter, then evaluated the difference in performance between the two models (ANOVA function of R) obtaining a p-value P_anova_real_<2.2e-16. Since the different observations are not independent, the analytically estimated P_anova_real_ considers inflated degrees of freedom and is not reliable. To obtain a statistically correct estimation of the difference in performance between the full and nested model, we performed a randomisation test. We computed 10000 permutations of the AUC versus parameter association, and fitted the full and nested model on randomised data as before. Noticeably, randomised data maintained the correlation structure of real data and only the association between the AUC and the independent variables was lost. For each permutation we computed the p-value P_anova_random_ comparing full and nested models. To measure the relevance of the minimal length parameter, we counted the number of permutations resulting in P_anova_random_<=P_anova_real_. Since we did not obtain any random p-value less than the real one, we can say that the minimal length parameter is relevant and we can estimate its probability as less than 1e-4.

Subsequently, we considered the full parameters space of 3plex (formula: AUC ∼ peak filtering method + ssRNA (or technique) + n peaks + single strandedness + minimal length + error rate + guanine rate + repeat filter + consecutive errors + scoring method), still focusing on the 7 lncRNA having experimental evidence of functional TPX. To reduce the computational time we considered only the values 8 and 10 for the minimal length. We performed a permutation test as previously explained for each parameter and we obtained a p-value lower than 1e-4 for all of them except for minimal length (p-value < 1e-3) and consecutive error (not significant).

To efficiently explore the parameters space considered we leveraged the Paramspace functionality of snakemake (27).

### 3.7. TF binding sites prediction evaluation

A set of TFs for binding sites prediction evaluation was selected from Jayaram et al. (28) in the human K562 cell line (CTCF, E2F1, GATA2, IRF1, MAX, NFYA, TAL1, YY1). We downloaded the ChIP-seq IDR conservative thresholded peaks from the ENCODE database as true positive binding sites. Hence, we selected genomic random regions as negative controls, maintaining the same length distribution of the positive ones using bedtools shuffle as for lncRNAs binding sites. We downloaded the PWM models of binding sites for those TFs from JASPAR (29) (respectively MA0139, MA0024, MA0036, MA0050, MA0058, MA0060, MA0091, MA0095) and used FIMO (30) from the MEME suite (31) (one of the most used TF motif scanning tools) with standard parameters in order to associate a score to each genomic region. The obtained values were employed to evaluate the performance of the TFs binding sites predictor using ROC and AUC values, computed with the pROC r package (25) as for lncRNAs binding sites.

## 4. Results

### 4.1. RNABSdb

We surveyed the literature and found experimentally determined binding sites for different lncRNAs in human, mouse, and Drosophila (see Supplementary Table 1 for details).

Data obtained employing RNABS-seq techniques are frequently processed using custom bioinformatic pipelines and published as a list of binding sites. Coming from different laboratories, these lists can be biassed by custom parameter settings, cutoffs and different software used.

For this reason, we collected and re-analyzed the raw-data in a uniform way for 16 human and 12 murine lncRNAs for which raw data were available.

Previous studies showed that, in contrast to histone modifications, which often broadly occupy genomic regulatory elements such as promoters and enhancers, lncRNAs occupancy tends to be focal, interspersed, and gene-selective in nature, more resembling the transcription factors pattern (32).

We employed the ENCODE transcription factor ChIP-seq data processing pipeline that is suitable for molecules that are expected to bind DNA in a punctate manner (Figure 1).

For this analysis, we focused on ChIRP-seq, CHART-seq and ChOP-seq data that allow the identification of individual lncRNA and we excluded ChAR-seq, GRID-seq and MARGI-seq data.

To enable systematic studies on transcriptional regulation driven by lncRNAs we made the harmonised data available through the RNABSdb website. RNABSdb allows the user to submit new data that will also be collected in the database upon proper revision. Moreover, we provide, as a service, the analysis of unpublished raw data with the pipeline we describe in this paper, thus allowing the proper comparison of new data with all the collected references.

RNABS-seq data analysis and triplex prediction depend on the choice of many different parameters, among which some are more relevant than others. We chose to deeply investigate the parameter space of triplex forming prediction algorithms as they have been less extensively tested with respect to ChIP-seq analysis pipelines.

For the RNABS-seq techniques, we decided to focus on the strategy used to handle replicates, namely 1) IDR-conservative, 2) IDR-optimal, 3) Overlap-coservative, 4) Overlap-optimal, 5) the top 1000 IDR or Overlap peaks (see methods) and 6) the set of peaks provided by the authors as bed file if available. This parameter is indeed the one with the stronger impact on the final number of called peaks. For sake of simplicity we refer to these different strategies as “peak filtering methods’’. The Irreproducible Discovery Rate (IDR) is the approach suggested by ENCODE and measures the reproducibility of an assay (23).

The IDR-conservative method applies the IDR procedure on the true replicates (in RNABS-seq typically arising from even and odd probes), whilst IDR-optimal considers pseudoreplicates too, including those obtained by subsampling pooled replicates. The optimal set is more sensitive, in particular when one of the replicates has substantially lower data quality than the other. The Overlap peaks set is obtained considering overlapping peaks employing bedtools intersect; also in this case conservative sets consider true replicates while optimal sets consider pseudoreplicates.

By applying the ENCODE pipeline and varying this parameter we determined the number of binding sites for each lncRNA. Starting from IDR-conservative filtered peaks we observed that only 16 out of 33 lncRNAs show at least 100 binding sites, but this number increases to 23 when considering IDR optimal filtered peaks (Figure 2). Moreover, in many cases we observed 3 orders of magnitude more peaks in the optimal version compared with the conservative one, impling poor reproducibility, maybe due to poor quality of one of the replicates. According to ENCODE recommendations (see methods), about one third of experiments reach an “ideal” reproducibility rate. Focusing on this high quality subset, we noticed a difference in IDR-conservative peaks numbers ranging from 649 (mouse DT926623.1) to 75109 (human HAND2-AS1) (Figure 2 and Supplementary Table 2).

**Figure 2.**
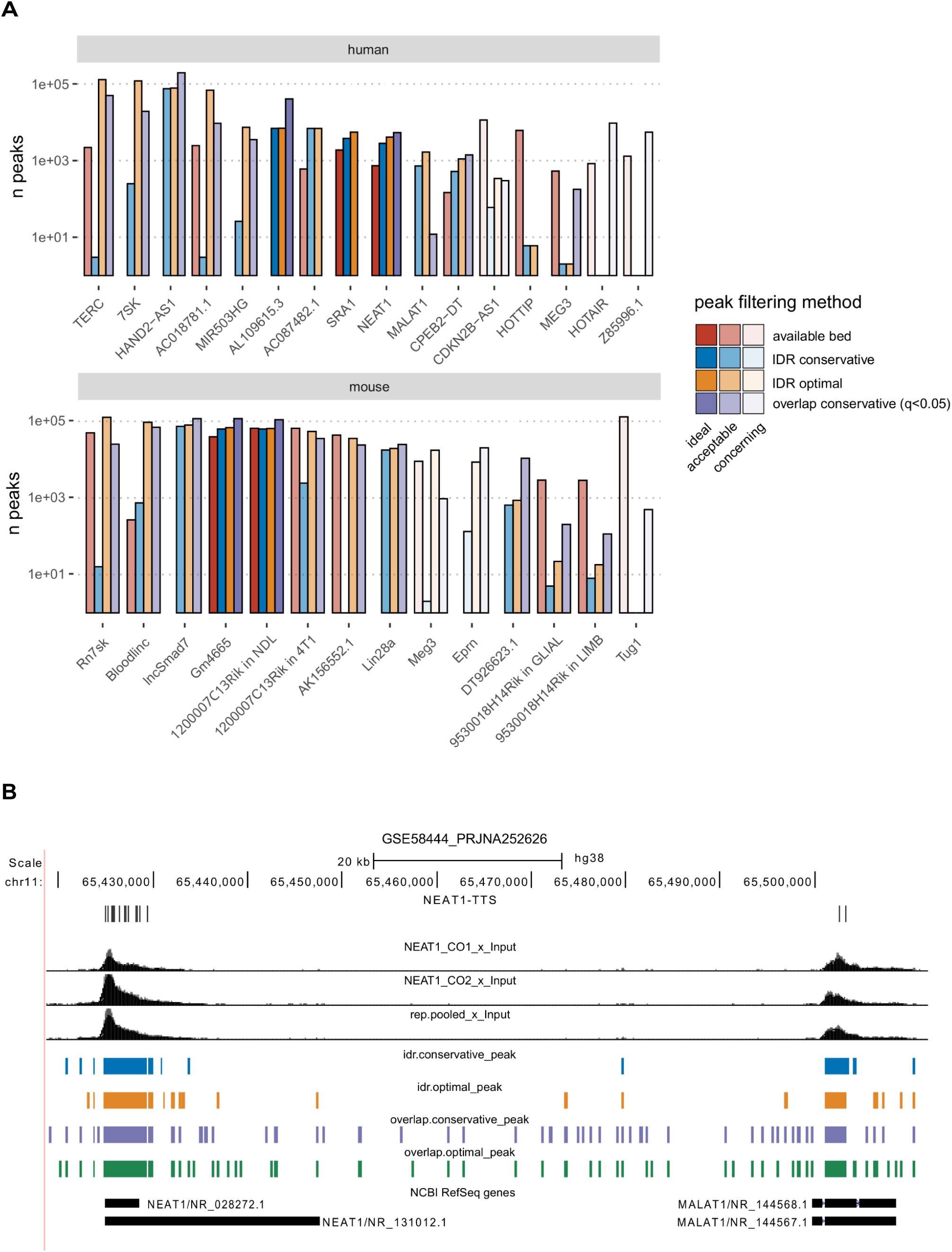
(**A**) DNA Binding sites count of lncRNAs using different peak filtering methods in human and mouse. Different colours discriminate among the customised called peaks produced by the authors (red), IDR conservative consistent peaks (blue), IDR optimal consistent peaks (yellow) and q-value filtered Overlap conservative consistent peaks (violet). Three different transparency levels are used to represent the replicate concordance: dark colours when “ideal”, medium transparency when “acceptable” and high transparency when “concerning”. The data are available in Supplementary Table 2. (**B**) Genome Browser view of the hg38 region involving NEAT1 and MALAT1 genes showing NEAT1 CHART-seq signals and peaks called using different peak filtering methods. The TTSs predicted by 3plex for NEAT1 on the experimental binding sites are shown in grey on the top.

At first, we asked if the great variability in the number of peaks depends on quality control (QC) parameters (Supplementary Table 3). Interestingly, we found that the number of IDR conservative peaks does not correlate significantly with any QC metrics (Spearman correlation test). This reassures us that the bona-fide number of peaks found with this method (theoretically more stringent than others) does not critically depend on technical variability among experiments, but possibly reflects intrinsic biological characteristics of different lncRNAs.

With this initial analysis we could establish that a typical DNA-binding lncRNA displays hundreds to tens of thousands DNA binding sites when the RNABS-seq is consistent among replicates.

Other methods often recall more peaks than IDR conservative, but are affected by technical variability (measured with different QC metrics),including the number of uniquely mapped reads, normalised strand cross-correlation, library complexity and Jensen-Shannon distance in fingerprint (Supplementary Table 4).

The available data allowed us to identify differences in the number of peaks determined using ChIRP-seq, CHART-seq or ChOP-seq, but, at this stage, the statistics are too poor to draw conclusions about the accuracy of different techniques.

### 4.2. 3plex implementation

We developed 3plex, a software tool to predict the DNA:DNA:RNA TPX. 3plex is built on top of Triplexator and implements further TPX evaluation methods based on thermal stability derived by LongTarget (17) and RNA secondary structure prediction.

The input of the 3plex algorithm is a single stranded RNA sequence (ssRNA) and a set of double stranded DNA sequences (dsDNAs). The output consists of 1) all possible TPXs that satisfy a set of constraints derived from Triplexator with associated thermal stability evaluation, and 2) a score for each dsDNA that predicts the capability of ssRNA to bind to it.

### 4.3. 3plex and RNABS-seq techniques evaluation

Among lncRNAs for which DNA binding sites have been determined, we identified 7 (TERC, NEAT1, HOTAIR, MEG3, ANRIL, Meg3, lncSmad7) for which the TPX binding mechanism have been experimentally validated (Supplementary Table 1). For these 7 lncRNAs that are known to form triple helices, we expected a good fraction of RNABS with high predicted triplex score. So we could use experimentally identified lncRNA binding sites as a benchmark to evaluate the goodness of triplex prediction approaches. Noteworthy, we could also do the other way round: we could evaluate the different RNABS-seq analysis parameters according to our TPX prediction.

To evaluate the concordance between experimental RNABS and predicted TTS we used the Area Under the Receiver Operator Curve (AUC). For each lncRNA we obtained a different RNABS set by varying the definition of true binding sites (changing the replicate handling strategy of ENCODE pipeline) and using different TTSs sets (by varying many 3plex parameters). Each combination of binding-sites and 3plex parameters produces a different AUC, so we used multivariate linear regression analysis to estimate the relevance of each parameter. Since different instances of parameters are clearly non independent, the statistical estimation was made using permutation tests that preserve the interdependencies in the null hypothesis.

Then we included in our analysis also other lncRNAs in RNABSdb, not having strong experimental evidence of functional TPX. With the same method, we discovered statistical evidence supporting the hypothesis that many of these lncRNAs directly bind the DNA via triplex formation.

### 4.3.1. Minimal TPX length and error rate

In order to investigate the influence of the minimal TPX length on the prediction, we chose to compare Triplexator default minimal TPX length of 16 with shorter lengths. Indeed, experimental triplex evidence revealed functional TPXs of few nucleotides (see Introduction). For this analysis, we only took into consideration the 7 lncRNAs with reported evidence of functional TPXs. We fixed Triplexator parameters to the values suggested by Matveishina et al. (26) and excluded 3plex specific parameters (thermal stability and secondary structure). To evaluate the relevance of minimal TPX length we first computed AUC values for minimal length 8, 10, 12 and 16, and then we compared a multivariate linear model that investigates the dependency of AUC on different parameters (minimal TPX length, number of peaks, peak filtering method and different ssRNA) with a similar nested model where the dependency on minimal TPX length is removed. We observed a significant influence of this parameter (p-value < 1e-04, permutation test), in particular the best minimal TPX length values were 8 and 10 (equally). The default cutoff of 16 resulted to be the worst, producing an average reduction in AUC of 0.1, while 12 gave an intermediate performance (Figure 3).

**Figure 3.**
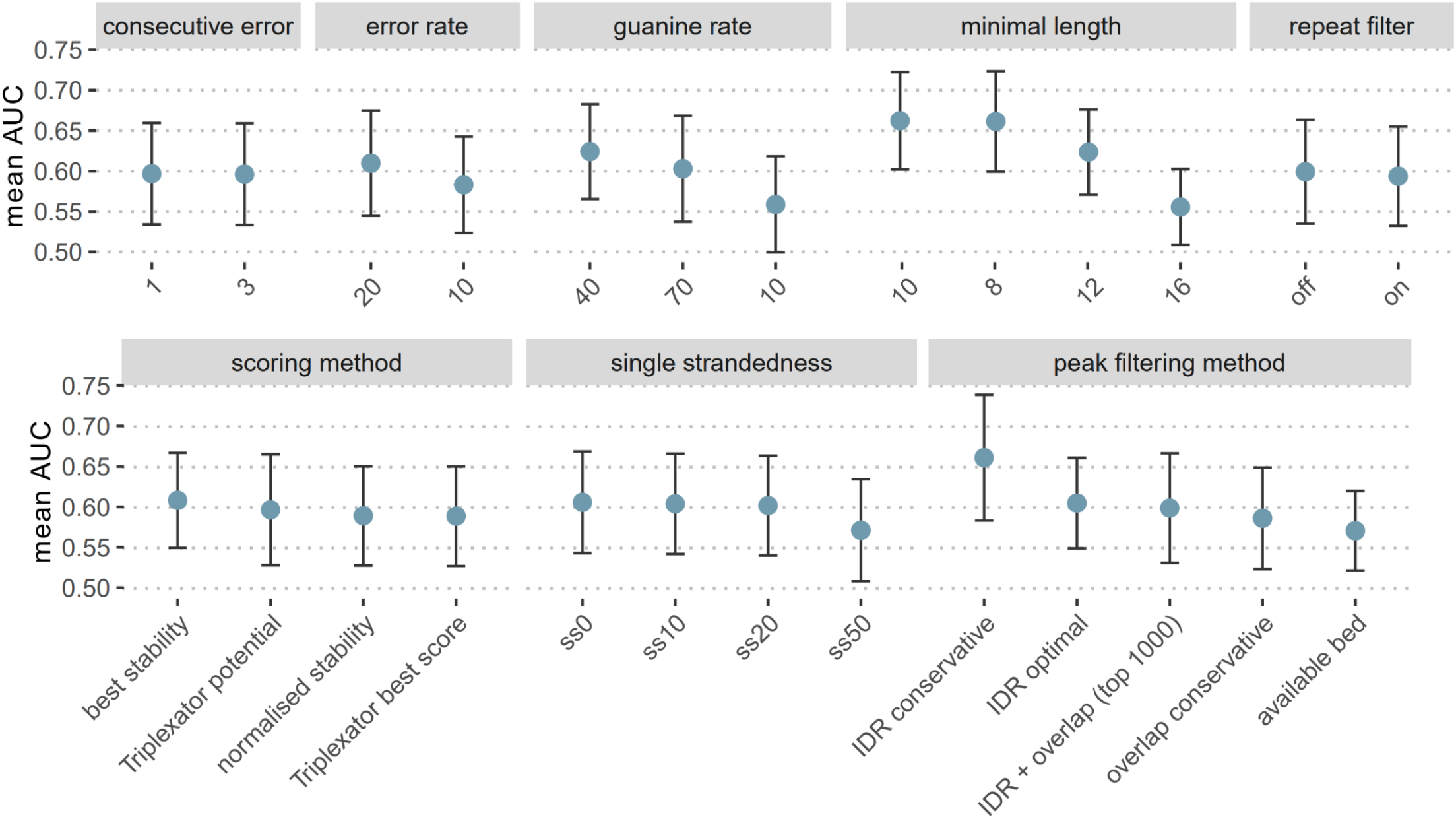
Impact of various parameters in triplex prediction accuracy. 3plex prediction performance is expressed as AUC obtained by comparing triplex scores of RNA binding sites in RNABSdb with randomised negative controls. Every parameter combination tested produces an AUC value for each lncRNA. For all the levels of the investigated parameters, the average AUC is represented as a blue dot and the standard deviation as error bars. The values for the minimal length are computed considering only a subset of the parameter space (see Methods).

Thus, we fixed the minimal TPX length to 8 or 10 and further investigated 3plex parameters space by introducing other parameters in the linear model (single strandedness cutoff, minimal error rate, guanine rate, repeat filter, maximal number of consecutive error, scoring method). Again, we compared the dependency of the AUC from these parameters with a similar nested model where the dependency from minimal TPX length is removed. We obtained a moderate, but significant preference for a value of 8 over 10 (average AUC difference of 0.005, p-value < 9e-4, permutation test).

It’s to be noted that 8 is below the minimum available threshold of Triplexator (thus we modified the source code), meanwhile LongTarget only focuses on long TPX.

Our observation highlights the fact that typical TPX are shorter than previously expected. Because of the long computational time required, we did not extensively test lower values for this parameter, but we evaluated minimal TPX length of 6 in a subsample, confirming that the signal to noise ratio was too poor and not obtaining any further improvement.

With the same strategy used to evaluate the relevance of the minimal TPX length, we also evaluated the relevance of the error rate. In this context, an error is a non-proper pairing in the TTS and TFO that forms a TPX, as well as deviation from the rules that sequences forming TSSs and TFOs should satisfy. The linear model shows that this parameter is relevant (p-value < 1e-4 permutation test). Setting the error rate to 20% resulted in an increase of average AUC of 0.02 compared to 10%. Moreover, we tested if the number of consecutive errors was relevant and we observed no significant differences in average AUC, allowing for just 1 or up to 3 consecutive errors.

These data suggest that TPXs are generally relatively small and degenerated. This observation could be surprising, since it seems that such models account for too little information to be biologically relevant. Actually these binding models do not differ much from many famous TF binding models (see Methods Section 4.7), considering their typical length and degeneration. This is reflected by the comparison of ROC curves describing the prediction of RNA binding sites and those describing the prediction of TF binding sites (Figure 4).

**Figure 4.**
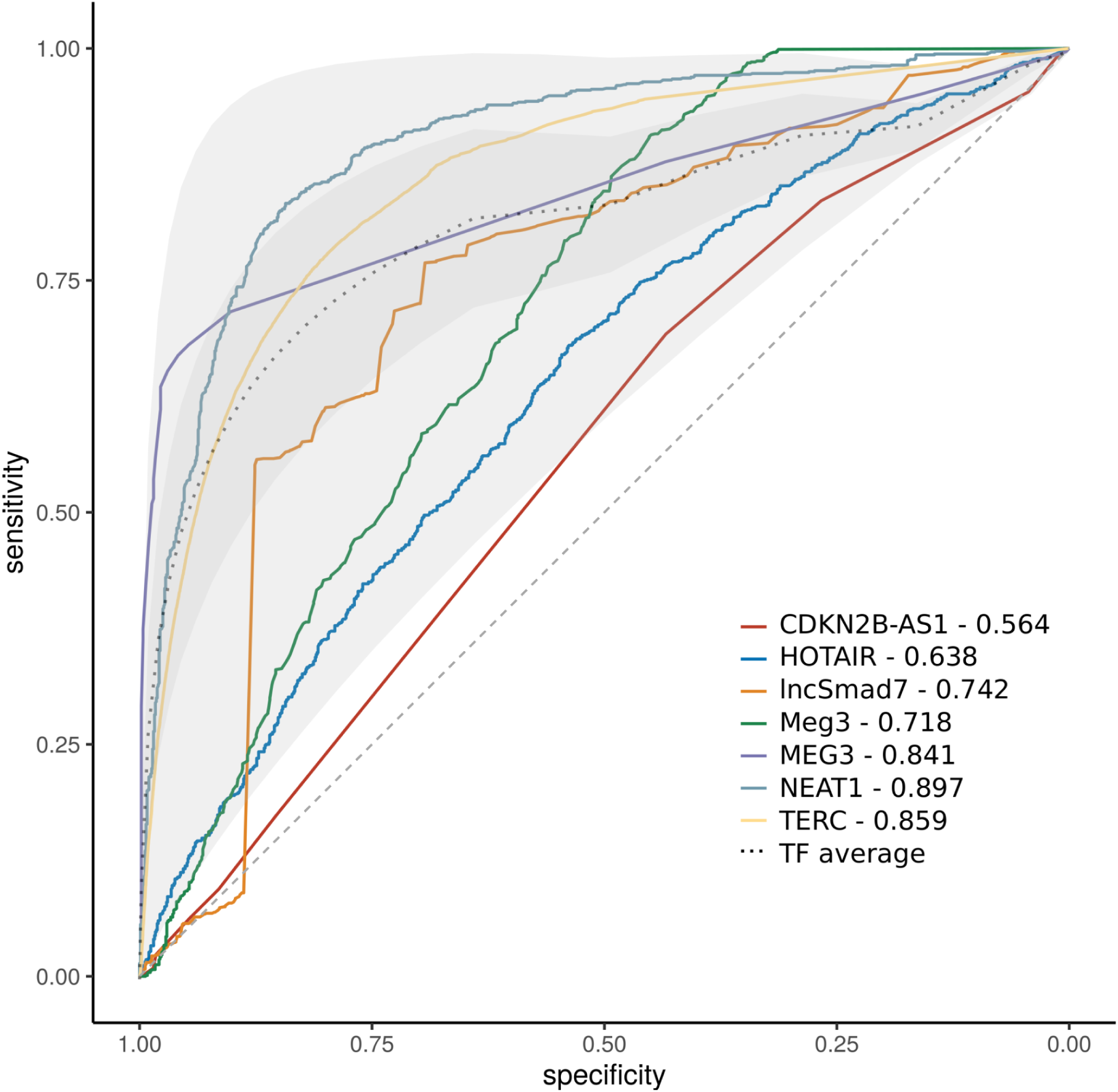
ROC curves showing the performance of lncRNAs TPX prediction compared with TFs binding sites prediction. ROC curves for each lncRNA with experimental evidence of functional TPX considering the 3plex parameter combination that produced the best AUC value (reported in the figure legend) (see Supplementary Table 6). The grey dashed line represents the performance of a random classifier. The grey dotted line illustrates the average ROC curve computed from the performance evaluation of FIMO TFs binding sites predictions. The dark grey shade represents the standard deviation from the average ROC curves for TFs, the light grey shade boundaries correspond to the worst and best TF ROC curves.

### 4.3.2. Effect of the thermal stability computation

3plex implements a thermal stability evaluation of TPX similar to the one developed in LongTarget (17). In particular, given a ssRNA and a dsDNA, 3plex finds all TPXs that satisfy the constraints as in Triplexator, then reports the best and the average stability over all TPXs. To evaluate the probability that a lncRNA may form a triplex on a given dsDNA sequence, we compared 4 alternative triplex scoring methods:

1. The Triplexator potential (the commonly used triplex potential returned by Triplexator, that is the length normalised number of TPXs (15)).
2. The normalised stability (the length normalised average thermal stability of TPXs).
3. The best stability (the stability score of the more stable among predicted TPXs).
4. The Triplexator best score (the highest Tripelxator score among predicted TPX, where the score is computed as the TPX length minus errors, such as mismatches or nucleotides not suitable for the TFO and TTS model).

Focusing on the 7 lncRNAs with reported evidence of functional TPXs, we investigated the dependency of AUC on different parameters. We considered a full multivariate linear model including all studied parameters as covariates (Peak_filtering_method, ssRNA, n_peaks, singleStrandedness, minimal_length, error_rate, guanine_rate, repeat_filter, consecutive_error, triplex_scoring_method) and evaluated the effect of the triplex scoring method by comparing the full model with a similar nested model where the dependency from triplex scoring method is removed. We found that the AUC depends on this parameter (p-value < 1e-4, permutation test), in particular the best stability method scored as best and produced an increase in average AUC of 0.01, if compared with the usual triplex potential (Figure 3).

Considering each lncRNA individually, the parameters sets that produce the best AUC require a stability based scoring method (best stability or normalised stability) for 4 out of 7 lncRNAs. Changing the scoring method and maintaining fixed all the other parameters we found that there was a statistically significant reduction in AUC when the scoring method was not based on stability (best Triplexator score or Triplexator potential). On the contrary, 3 lncRNAs show the best AUC with Triplexator score or Triplexator potential but in this case the difference in AUC with respect to the one computed using a stability based scoring method is not significant (see Supplementary Table 5).

### 4.3.3. Peak filtering method

The use of different peak filtering methods (described in section 3.1) is one of the covariates included in the multivariate model. By comparing the full model with the corresponding nested one and excluding this parameter, we found that it is relevant for the AUC (p-value < 1e4, permutation test). Interestingly, we observed that the RNABS redefined using the ENCODE pipeline showed on average higher AUC value than the sets of peaks provided by the authors. The best method was the IDR conservative one, producing an increase in average AUC of 0.09 compared with published peaks sets. Only 2 out of 7 lncRNA considered in this analysis had a defined set of IDR conservative peaks, while the other 5 showed insufficient reproducibility among replicates to apply this method. Anyway, we controlled for different lncRNAs as covariate in our model. Moreover, considering IDR optimal peaks that are available for 5 out of 7 lncRNA, we still obtained a significant increase in average AUC of 0.015 (Fig 3).

This highlights the importance of uniform and standard processing of RNABS-seq data and the relevance of the proposed RNABSdb, while the peaks derived from custom analysis by the authors of single papers (available for 6 out of 7 lncRNAs) showed the worst performance.

### 4.3.4. RNA secondary structure

In order to establish Hoogsteen hydrogen bonds with a DNA double helix, the nucleotides of the RNA molecule need to be in a free conformation. Previous studies report improvements in triplex prediction of specific lncRNAs by integrating RNA secondary structure information (Matveishina et al., 2020), but none of the triplex prediction software available takes into account this information. In 3plex we implemented a Single Strandedness (ss) parameter that determines the importance of secondary structure in the prediction of TPX, by masking the RNA sequence according to the probability of base pairing (see Methods). Despite testing different extent of sequence masking (0%, 10%, 20% or 50%), we found that, on average, the best AUC value is obtained by keeping the RNA sequence completely unmasked. Surprisingly, this means that the secondary structure of RNA does not affect TPX prediction in general and that TFO can be formed also in lncRNA regions likely to be in a double strand conformation.

Nevertheless, looking at each individual experiment one by one, and considering the full set of lncRNAs (including those yet lacking strong experimental evidence of functional TPX), 10 cases show improvement in AUC using secondary structure filtering. Notably, Lin28a AUC reaches 0.60 without filtering and 0.95 masking the 10% of most probable paired nucleotides in the RNA, highlighting the relevance of this 3plex feature in some cases. Further studies would be required to identify eventual secondary structure characteristics typical of these lncRNAs.

### 4.3.5. Other parameters

According to constraints due to Hoogsteen hydrogen bonding rules, a minimum guanine rate in the TTS is often required in TPX prediction. Our analysis shows that this parameter is relevant (p-value <1e-4, permutation test) and in general a minimum rate of 40% is preferable with respect to 70% or 10% and produces an increase in average AUC of 0.06 compared to 10%.

The option of filtering the repeats on the DNA sequences is available in Triplexator. The choice of this parameter significantly impacts the AUC in general (p-value <1e-4, permutation test) and avoiding filtering improves the triplex predictions.

It would be interesting to evaluate if different RNABS-seq techniques produce different results in terms of TPX AUC. Unfortunately, the data available are not sufficient to investigate this aspect.

### 4.4. Evidence of TPX potential for many lncRNA

In RNABSdb we have 20 human and murine lncRNAs with publicly available RNABS-seq data, but without experimental evidence of functional TPX. For two of them, the authors tested the capability of forming TPXs in silico (AC087482.1, Eprn) (10,33) and for Tug1 (34), the TPX mechanism of binding was only speculated.

We wondered for how many lncRNAs we could observe statistical evidence supporting a mechanism of DNA binding involving TPX formation. We used 3plex to predict TPX on the available binding sites collected in RNABSdb. We considered those sites as positive controls and created a randomised selection of genomic regions as negative control (See Methods), that we used to investigate the same parameter space as before.

Interestingly, we found 4 lncRNAs having an AUC above 0.8 (Lin28a, AK156552.1, HAND2-AS1, MALAT1), 4 lncRNAs above 0.7 (AL109615.3, 7SK, 1200007C13Rik, CPEB2-DT) and 6 lncRNAs above 0.6 (Rn7sk, AC087482.1, Tug1, Eprn, AC018781.1, 1200007C13Rik@NDL, SRA1, MIR503HG, Bloodlinc). All the mentioned AUC values have a corresponding highly-significant p-value, even if we consider the high number of tests due to the parameter space exploration (FDR < e-10, Mann-Whitney test with Benjamini-Hochberg correction), suggesting that they may bind DNA forming TPXs (Figure 5, supplementary table 6).

**Figure 5.**
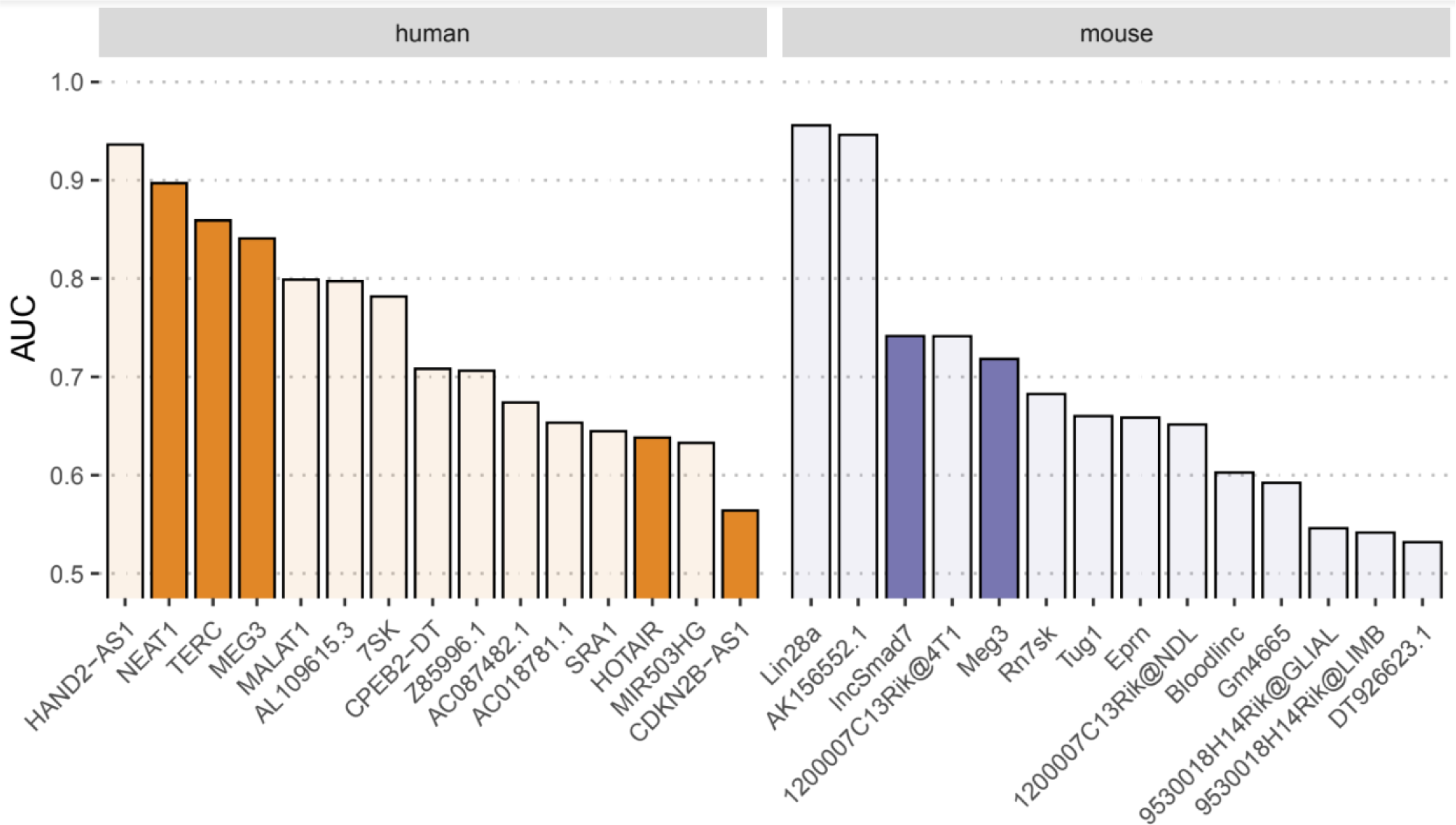
AUC values of all the analysed lncRNAs. The AUC values are reported from the best parameter set for each lncRNA (see Supplementary Table 6). Darker bars highlight lncRNAs with experimental TPX evidence.

## 5. Discussion

RNABSdb collects high throughput data regarding the binding of lncRNA on the human and murine genome. By uniform re-analysis of available raw data using the ENCODE pipeline, we observe a range in experimental quality. We estimated that a typical DNA-binding lncRNA can have from hundred to tens of thousands binding sites, and we argued that this quantification corresponds to a biological property of specific lncRNAs rather than to technicalities of experiments. This is because the number of binding sites does not significantly depend upon quality control parameters (e.g. number of mapped reads), as long as the replicates are consistent with one another.

We applied a model of RNA:DNA:DNA triple helices implemented in 3plex to the collected binding sites and showed that lncRNAs tend to form TPX that are short (around 8-10 nucleotides long) and degenerate (around 10%-20% of errors). This observation could be surprising, since it seems that such models account for too little information to be biologically relevant. Actually they do not differ much from that of typical DNA binding sites of many important transcription factors, considering their typical length and degeneration. This is reflected by the comparison of ROC curves describing the prediction of RNA binding sites and those describing the prediction of TF binding sites (Figure 4). Indeed the information content of a typical TFBS is paradoxically low when compared to the complex regulatory program they orchestrate in eukaryotic gene regulation programs (35). These observations suggest that lncRNAs (as well as TF) may not work alone, reinforcing the common idea that eukaryotic gene regulation is a complex process involving cooperation and competition of many different molecules. Another possibility is that multiple short TPX close to each other in the same genomic region act cooperatively to enhance the specificity of the binding as observed in the X chromosome inactivation by Xist (36).

Actually in vitro biochemical experiments show that even TXP as small as 12 nt can be functional (37), and in a previous work we experimentally validated functional specificity of some short TPXs formed by lncSmad7 (11). Moreover Matveishina et al elaborated a practical guidance in genome-wide RNA:DNA triple helix prediction using experimentally validated binding sites of 4 lncRNA and found that the minimal available cutoff in TPX length of Triplexator produced the best prediction(26). We confirmed these findings on a much wider dataset.

Moreover, investigating the TPX potential of 20 lncRNA not previously reported to bind DNA using this mechanism and having available binding sites in RNABSdb, we found strong evidence of TPX formation for 4 lncRNAs (Lin28a, AK156552.1, HAND2-AS1, MALAT1), good evidence for other 4 lncRNAs (AL109615.3, 7SK, 1200007C13Rik, CPEB2-DT), and moderate evidence for other 8 (Rn7sk, AC087482.1, Tug1, Eprn, AC018781.1, SRA1, MIR503HG, Bloodlinc) suggesting that DNA binding forming TPX could be a widespread mechanism adopted by lncRNAs. This idea was previously reported in literature (16,38) using purely computational methods. RNABSdb allows the first large-scale evaluation of this hypothesis leveraging high-throughput experimental data.

Hour work suggests guidelines to genome wide TPX prediction. Moreover, 3plex improves the state of the art in this field. Using RNABSdb data we showed the relevance of integrating Triplexator rules with thermal stability evaluation and discussed the impact of RNA secondary structure prediction.

## 6. Data availability

RNABSdb is available through https://molinerislab.github.io/RNABSdb/. For the sake of simplicity and maintainability all data are simply available as static downloadable files. Binding sites and signal levels are also visualizable on the UCSC genome browser (see figure 2B).

3plex source code is available at https://github.com/molinerisLab/3plex.

## 7. Acknowledgements

We thank Danny Incarnato, Department of Molecular Genetics, University of Groningen and Paolo Provero, Dipartimento di Neuroscienze, University of Torino for the helpful discussion of the results.

## Bibliography

1. Mattick JS. RNA regulation: a new genetics? Nat Rev Genet. 2004 Apr;5(4):316–23.

2. Liu SJ, Nowakowski TJ, Pollen AA, Lui JH, Horlbeck MA, Attenello FJ, et al. Single-cell analysis of long non-coding RNAs in the developing human neocortex. Genome Biol. 2016 Dec 14;17(1):67.

3. Wu H, Kirita Y, Donnelly EL, Humphreys BD. Advantages of Single-Nucleus over Single-Cell RNA Sequencing of Adult Kidney: Rare Cell Types and Novel Cell States Revealed in Fibrosis. J Am Soc Nephrol. 2019 Jan;30(1):23–32.

4. Alessio E, Bonadio RS, Buson L, Chemello F, Cagnin S. A Single Cell but Many Different Transcripts: A Journey into the World of Long Non-Coding RNAs. Int J Mol Sci. 2020 Jan 1;21(1):302.

5. Pinkney HR, Black MA, Diermeier SD. Single-Cell RNA-Seq Reveals Heterogeneous lncRNA Expression in Xenografted Triple-Negative Breast Cancer Cells. Biology. 2021 Sep 30;10(10):987.

6. Jiang C, Li Y, Zhao Z, Lu J, Chen H, Ding N, et al. Identifying and functionally characterizing tissue-specific and ubiquitously expressed human lncRNAs. Oncotarget. 2016 Feb 9;7(6):7120–33.

7. Statello L, Guo CJ, Chen LL, Huarte M. Gene regulation by long non-coding RNAs and its biological functions. Nat Rev Mol Cell Biol. 2021 Feb;22(2):96–118.

8. Sun JS, Hélène C. Oligonucleotide-directed triple-helix formation. Curr Opin Struct Biol. 1993 Jun 1;3(3):345–56.

9. Duca M, Vekhoff P, Oussedik K, Halby L, Arimondo PB. The triple helix: 50 years later, the outcome. Nucleic Acids Res. 2008;36(16):5123–38.

10. Zapparoli E, Briata P, Rossi M, Brondolo L, Bucci G, Gherzi R. Comprehensive multi-omics analysis uncovers a group of TGF-β-regulated genes among lncRNA EPR direct transcriptional targets. Nucleic Acids Res. 2020 Sep 18;48(16):9053–66.

11. Maldotti M, Lauria A, Anselmi F, Molineris I, Tamburrini A, Meng G, et al. The acetyltransferase p300 is recruited in trans to multiple enhancer sites by lncSmad7. Nucleic Acids Res. 2022 Mar 21;50(5):2587–602.

12. Brown JA. Unraveling the structure and biological functions of RNA triple helices. Wiley Interdiscip Rev RNA. 2020;11(6):1–26.

13. Li Y, Syed J, Sugiyama H. RNA-DNA Triplex Formation by Long Noncoding RNAs. Cell Chem Biol. 2016;23(11):1325–33.

14. Kuo CC, Hänzelmann S, Sentürk Cetin N, Frank S, Zajzon B, Derks JP, et al. Detection of RNA-DNA binding sites in long noncoding RNAs. Nucleic Acids Res. 2019;47(6):e32.

15. Buske FA, Bauer DC, Mattick JS, Bailey TL. Triplexator: Detecting nucleic acid triple helices in genomic and transcriptomic data. Genome Res. 2012;22(7):1372–81.

16. Jalali S, Singh A, Maiti S, Scaria V. Genome-wide computational analysis of potential long noncoding RNA mediated DNA: DNA: RNA triplexes in the human genome. J Transl Med. 2017;15(1):1–17.

17. He S, Zhang H, Liu H, Zhu H. LongTarget: A tool to predict lncRNA DNA-binding motifs and binding sites via Hoogsteen base-pairing analysis. Bioinformatics. 2015;31(2):178–86.

18. Engreitz JM, Pandya-Jones A, McDonel P, Shishkin A, Sirokman K, Surka C, et al. The Xist lncRNA Exploits Three-Dimensional Genome Architecture to Spread Across the X Chromosome. Science. 2013 Aug 16;341(6147):1237973.

19. Bezzecchi E, Pagani G, Forte B, Percio S, Zaffaroni N, Dolfini D, et al. MIR205HG/LEADR Long Noncoding RNA Binds to Primed Proximal Regulatory Regions in Prostate Basal Cells Through a Triplex- and Alu-Mediated Mechanism. Front Cell Dev Biol. 2022 Jun 17;10:909097.

20. Ducoli L, Agrawal S, Sibler E, Kouno T, Tacconi C, Hon CC, et al. LETR1 is a lymphatic endothelial-specific lncRNA governing cell proliferation and migration through KLF4 and SEMA3C. Nat Commun. 2021 Dec;12(1):925.

21. Landt S, Marinov G. ChIP-seq guidelines and practices of the ENCODE and modENCODE consortia. Genome &. 2012;(Park 2009):1813–31.

22. Kharchenko PV, Tolstorukov MY, Park PJ. Design and analysis of ChIP-seq experiments for DNA-binding proteins. Nat Biotechnol. 2008 Dec;26(12):1351–9.

23. Li Q, Brown JB, Huang H, Bickel PJ. Measuring reproducibility of high-throughput experiments. Ann Appl Stat. 2011;5(3):1752–79.

24. Lorenz R, Bernhart SH, Höner zu Siederdissen C, Tafer H, Flamm C, Stadler PF, et al. ViennaRNA Package 2.0. Algorithms Mol Biol. 2011 Dec;6(1):26.

25. Robin X, Turck N, Hainard A, Tiberti N, Lisacek F, Sanchez JC, et al. pROC: an open-source package for R and S+ to analyze and compare ROC curves. BMC Bioinformatics. 2011 Jan;12(1):77.

26. Matveishina E, Antonov I, Medvedeva YA. Practical guidance in genome-wide RNA: DNA triple helix prediction. Int J Mol Sci. 2020;21(3).

27. Köster J, Rahmann S. Snakemake-a scalable bioinformatics workflow engine. Bioinformatics. 2012;28(19):2520–2.

28. Jayaram N, Usvyat D, Martin AC. Evaluating tools for transcription factor binding site prediction. BMC Bioinformatics. 2016;17(1):1–12.

29. Castro-Mondragon JA, Riudavets-Puig R, Rauluseviciute I, Lemma RB, Turchi L, Blanc-Mathieu R, et al. JASPAR 2022: the 9th release of the open-access database of transcription factor binding profiles. Nucleic Acids Res. 2022 Jan 7;50(D1):D165–73.

30. Grant CE, Bailey TL, Noble WS. FIMO: scanning for occurrences of a given motif. Bioinformatics. 2011 Apr 1;27(7):1017–8.

31. Bailey TL, Boden M, Buske FA, Frith M, Grant CE, Clementi L, et al. MEME SUITE: tools for motif discovery and searching. Nucleic Acids Res. 2009 Jul 1;37(Web Server issue):W202–8.

32. Chu C, Qu K, Zhong FL, Artandi SE, Chang HY. Genomic Maps of Long Noncoding RNA Occupancy Reveal Principles of RNA-Chromatin Interactions. Mol Cell. 2011 Nov;44(4):667–78.

33. Merry CR, Forrest ME, Sabers JN, Beard L, Gao XH, Hatzoglou M, et al. DNMT1-associated long non-coding RNAs regulate global gene expression and DNA methylation in colon cancer. Hum Mol Genet. 2015 Nov 1;24(21):6240–53.

34. Long J, Badal SS, Ye Z, Wang Y, Ayanga BA, Galvan DL, et al. Long noncoding RNA Tug1 regulates mitochondrial bioenergetics in diabetic nephropathy. J Clin Invest. 2016 Oct 17;126(11):4205–18.

35. Kribelbauer JF, Rastogi C, Bussemaker HJ, Mann RS. Low-Affinity Binding Sites and the Transcription Factor Specificity Paradox in Eukaryotes. Annu Rev Cell Dev Biol. 2019 Oct 6;35:357–79.

36. Matsuno Y, Yamashita T, Wagatsuma M, Yamakage H. Convergence in LINE-1 nucleotide variations can benefit redundantly forming triplexes with lncRNA in mammalian X-chromosome inactivation. Mob DNA. 2019 Dec;10(1):33.

37. Mondal T, Subhash S, Vaid R, Enroth S, Uday S, Reinius B, et al. MEG3 long noncoding RNA regulates the TGF-β pathway genes through formation of RNA-DNA triplex structures. Nat Commun. 2015;6.

38. Soibam B. Super-lncRNAs: Identification of lncRNAs that target super-enhancers via RNA:DNA:DNA triplex formation. Rna. 2017;23(11):1729–42.

